# Repeat-specific functions for the C-terminal domain of RNA polymerase II in budding yeast

**DOI:** 10.1101/274274

**Authors:** Michael Babokhov, Mohammad M. Mosaheb, Richard W. Baker, Stephen M. Fuchs

## Abstract

The C-terminal domain (CTD) of the largest subunit of RNA polymerase II (RNAPII) is required to regulate transcription and to integrate it with other essential cellular processes. In the budding yeast *Saccharomyces cerevisiae*, the CTD of Rpb1p consists of 26 conserved heptad repeats that are post-translationally modified to orchestrate protein factor binding at different stages of the transcription cycle. A long-standing question in the study of the CTD is if there are any functional differences between the 26 repeats. In this study, we present evidence that repeats of identical sequence have different functions based on their position within the CTD. We assembled plasmids expressing Rpb1p with serine to alanine substitutions in three defined regions of the CTD and measured a range of phenotypes for yeast expressing these constructs. Mutations in the beginning and middle regions of the CTD had drastic, and region-specific effects, while mutating the distal region had no observable phenotype. Further mutational analysis determined that Ser5 within the first region of repeats was solely responsible for the observed growth differences and sequencing fast-growing suppressors allowed us to further define the functional regions of the CTD. This mutational analysis is consistent with current structural models for how the RNAPII holoenzyme and the CTD specifically would reside in complex with Mediator and establishes a foundation for studying regioselective binding along the repetitive RNAPII CTD.

## Introduction

RNA polymerase II (RNAPII) is a 12-subunit complex responsible for the transcription of all mRNA in eukaryotes. The largest subunit of RNAPII, Rpb1p, contains a conserved C-terminal domain (CTD) connected to the catalytic core by a flexible linker. The CTD is required for RNAPII activity *in vivo*, acting as a binding site for proteins involved in transcription initiation and other essential co-transcriptional processes (CORDEN 2013; EICK AND GEYER 2013). The coordination of all of these processes by the CTD throughout the transcriptional cycle continues to be a topic of intensive research. The CTD consists of tandemly repeating peptide units with the consensus sequence of Y_1_S_2_P_3_T_4_S_5_P_6_S_7_. Budding yeast Rpb1p contains 26 repeats that adhere closely to the consensus sequence while vertebrates have 52 repeats with the distal half containing many repeats degenerate at the Ser7 position. Protein factor binding to the CTD is mediated by extensive post-translational modification of the five hydroxylated amino acids and isomerization of the two prolines within each repeat (LEE AND GREENLEAF 1989; FEAVER *et al.* 1991; ZHANG AND CORDEN 1991; VALAY *et al.* 1995; FUCHS *et al.* 2009). Ser2 and Ser5 phosphorylation are the most commonly studied modifications and are evenly distributed across the CTD repeats (SUH *et al.* 2016). Dynamic patterns of CTD modifications are proposed to form a CTD code that directs progress through the transcription cycle.

Extensive research has uncovered many of the essential functional elements within the repetitive CTD. Mutations to the modifiable residues in the CTD cause drastic phenotypes, although the specific residue requirements can differ between organisms (HSIN *et al.* 2011; SCHWER AND SHUMAN 2011). Although the consensus sequence is a heptad repeat, the functional unit of the CTD is actually defined by two consecutive repeats that contain properly spaced Tyr1, Ser2 and Ser5 residues (STILLER AND COOK 2004; LIU *et al.* 2008). This functional unit is in agreement with structural studies of CTD binding factors that show surface interactions with two or more repeats (KUBICEK *et al.* 2012; ROBINSON *et al.* 2012). Furthermore, while the wildtype number of repeats is strongly selected for in nature (NONET AND YOUNG 1989), less than half of the total repeats appear to be required for normal growth (BARTOLOMEI *et al.* 1988; WEST AND CORDEN 1995; SCHNEIDER *et al.* 2010). These findings demonstrate that there are additional determinants of CTD function beyond just the linear sequence of heptad repeats.

The large number of CTD repeats, beyond those needed to support growth, raises the possibility of a division of function between the different repeats. For example, in mammalian cells, the elongation factor Spt6p requires the N-terminal half of the CTD (YOH *et al.* 2008) while splicing and 3’ processing require the C-terminal half (FONG AND BENTLEY 2001). These differences could be explained either by the different heptad sequences found in the two halves of the mammalian CTD (CORDEN 2013), the distance of the region from the core of the polymerase holoenzyme, or a combination of these considerations. Work in the budding yeast CTD similarly uncovered functional differences between the two halves of the CTD (WEST AND CORDEN 1995; WILCOX *et al.* 2004). However, unlike the mammalian CTD, the yeast CTD consists almost exclusively of consensus repeats. Therefore, in the absence of extensive sequence differences there must be additional determinants, such as distance from the holoenzyme, that lead to functional specialization in the budding yeast CTD.

Previously we developed a genetic system to investigate instability and repeat number in the budding yeast CTD (MORRILL *et al.* 2016). Here, we use this system to examine the effects of position specific repeat mutation on cellular survival and gene expression. We find that serine to alanine mutations within blocks of repeat units have profoundly different effects on cell survival and several other phenotypes (e.g. salt stress, inducible growth, and 6-azauracil sensitivity (POWELL AND REINES 1996)), dependent on their location within the CTD. In particular, we found that mutations within the middle third of the CTD resulted in generally poor growth whereas mutations to the first eight repeats had growth defects specific to inositol auxotrophy. In contrast, mutations in the last eight repeats had no discernable effect in any conditions tested. The repetitive coding sequence of the CTD makes it prone to spontaneous mutagenesis (MORRILL *et al.* 2016). We exploited this property to identify and analyze plasmid-based spontaneous suppressors that would bypass the poor growth of our CTD mutants. From these suppressors, we identified two discreet windows within the CTD that are required for viability in the presence and absence of inositol. Based on existing structural models of RNAPII and the Mediator complex, we propose that these regions are responsible for coordinating CTD interactions with Mediator.

## Materials and Methods

### Yeast Strains and Plasmids

Yeast strains were cultured in standard media and grown at 30° C except where otherwise noted. All of the reported strains are derivatives of GRY3019 (MATa *his3Δ, leu2Δ, lys2Δ, met15Δ, trp1Δ*::hisG, URA::CMV-tTA, kanRPtetO7-TATA-RPB1) provided by the Strathern lab (MALAGON *et al.* 2006) or from the yeast deletion collection (WINZELER *et al.* 1999). Gene tagging cassettes were created using PCR and integrated by homologous recombination (JANKE *et al.* 2004). The full set of strains used in this study is listed in Table S1. Selection was performed in synthetic complete (SC) media or plates lacking the appropriate amino acid for auxotrophic strains (ADAMS *et al.* 1997). Dominant drug resistance markers KanMX6, HphNT1 and NatNT2 were selected for using 50 μg/mL of geneticin (G418), hygromycin B and nourseothricin (ClonNAT), respectively. Ammonium sulfate was replaced with 1 g/L of monosodium glutamate as a nitrogen source whenever these drugs were used for selection in liquid media or plates. Region-specific mutants, and serine-specific CTD variant plasmids were made using the recursive directional ligation by plasmid reconstruction method (MCDANIEL *et al.* 2010). The construction of full length consensus and truncated CTD plasmids was described previously (MORRILL *et al.* 2016). To build CTD plasmids with repeat specific mutations, oligonucleotides that coded for two repeat blocks of the sequence (PTAPAYA)_2_ were recursively ligated together using two base pair overhangs until the desired CTD sequences were obtained. Similar oligonucleotides were used to create the serine-specific constructs (e.g. PTAPSYS for S5A). These CTD sequences were then cloned into pRPB1 using *Sac*1 and *Xma*1 restriction sites and verified by sequencing as described previously (MORRILL *et al.* 2016).

### Spotting Assays

Cells were grown overnight at 30° C in SC–LEU media and diluted to OD_600_ 0.2 in fresh SC–LEU in the morning. Yeast were allowed to divide at least two times (OD_600_ 0.8 – 1.0) before being collected and washed twice in sterile water. Cell number was estimated by spectrophotometry (OD_600_ = 1 ∼ 1x10^7^ cells/mL) and suspensions were transferred to a 96 well plate (250μL of ∼1x10^7^ cells/mL) and serially diluted 5-fold in sterile water. The dilutions were then spotted onto the appropriate plates using a 48-pin replicator. The plates were grown at 30° C (except where indicated) and photographed daily starting at two days. All spotting experiments were performed a minimum of three times from independent plasmid transformations and a representative image was selected to display.

### Western Blotting

Western analysis of the block mutants was performed as described previously (MORRILL *et al.* 2016) with the following changes. Proteins were separated on an 8% SDS-PAGE gel made with a standard 37.5:1 acrylamide:bis-acrylamide ratio and transferred to PVDF. Membranes were incubated with the primary antibodies raised against: Rpb1p (Y-80, Santa Cruz), phosphorylated Ser2 (a generous gift from Dirk Eick), phosporylated Ser5 (clone 3E8 from Active Motif) and G6PDH loading control (Sigma A9521).

### INO1 expression

RNA extracts to assay *INO1* expression were prepared by growing cultures at 30° C in SC–LEU media with 50 μg/mL doxycycline (DOX) (Alfar Aesar) to mid log phase (OD_600_0.6 – 0.8). Cells were harvested, washed twice in sterile water to remove any remaining inositol and resuspended in SC–LEU+DOX media that lacked inositol (SC–LEU– INO+DOX). Cultures were grown for two hours without inositol to induce *INO1* gene expression. After induction, cells were harvested, washed and stored at -80° C. Total RNA was extracted using an Illustra RNAspin Mini kit (GE Healthcare), following the manufacturers protocol for yeast. RNA extracts were quantified using a NanoDrop 2000 spectrophotometer (Thermo Scientific) and 50 ng of total RNA was primed with poly-(dT) primers to obtain cDNA with a SuperScript First-Strand Synthesis kit (Invitrogen). One microliter of cDNA was used as a template to amplify the *INO1* gene and the resulting bands were quantified in ImageJ and normalized to *ACT1* levels. RT-PCR was performed with RNA from three independent cultures for controls and six independent cultures for –INO+DOX experimental samples. The mean is reported in the text and a two-way ANOVA was performed to assess significance using Graphpad Prism software. *Suppressor mutant screen*

Spontaneous suppressor mutations in block mutant plasmids were obtained using cultures grown in a 96 well plate. Individual colonies were taken from a fresh plasmid transformation and suspended in 1 mL of SC–LEU media in a deep well plate. Plates were grown at 30° C with occasional shaking for one day. The cultures were diluted to an OD_600_ 0.8 – 1.0 using fresh SC–LEU media and plated to SC–LEU plates with and without DOX and inositol using a 48-pin replicator. Plates were incubated at 30° C between three and five days until fast-growing colonies appeared. Colonies were screened by PCR with primers flanking the CTD coding region and loaded on a 1% agarose gel to identify plasmid-based mutational events as indicated by a change in band size relative to the amplified genomic *RPB1* CTD coding region. Mutated plasmids were extracted, sequenced and retransformed into GRY3019 to confirm their ability to support growth on media containing DOX.

### Structural Modeling

A cryoEM structure of the full PIC-Med complex from *S. cerevisiae* was recently determined to a resolution of ∼20 Å ((ROBINSON *et al.* 2016), EMD-8308). This structure includes RNAPII, TFIIA, TFIIB, TFIID, TFIIE, TFIIF, TFIIH, and TFIIS, and the full Mediator complex, including the Head, Middle, and Tail modules. Robinson et al. were able to make a molecular model for RNAPII, Mediator Head and Middle, and subsets of the general TFs (ROBINSON *et al.* 2012). To build on this model, we aligned the molecular model for the complete human TFIIH from a recent high resolution cryoEM structure (5of4.pdb) (GREBER *et al.* 2017). TFIIH was aligned using XPD and CXPD, the human homologs of Rad25 and Rad3. We also aligned a crystal structure of *Schizosaccharomyces pombe* RNAPII, which includes around 30 amino acids of the Rpb1p linker that makes contacts with the RNAPII subunits Rvb1p and Rvb7p ((SPAHR *et al.* 2009), 3h0g.pdb). All docking, alignment, and figure making was done using UCSF Chimera (PETTERSEN *et al.* 2004) or PyMol (The PyMol Molecular Graphics System, Version 1.7, Schrodinger, LLC).

### Reagent and Data Availability

The complete list of plasmids used in this is found in Table S1 and pertinent plasmids have been deposited to Addgene. Additional data regarding the CTD constructs are presented in Figure S1–S4. All reagents and data are available upon request.

## Results

### Repeat number requirements in phenotypes related to CTD function

Previous studies of RNAPII examined a number of phenotypes to dissect CTD repeat function (NONET *et al.* 1987; ARCHAMBAULT *et al.* 1996). We first determined the phenotypic consequences of varying CTD repeat number requirements in our TET-off system. Briefly, the addition of the antibiotic doxycycline (DOX) to growth media prevents expression of the genomic wildtype copy of *RPB1*. This leaves the plasmid-based copy containing our CTD constructs as the only source of Rpb1p for the cell. We tested a series of CTD truncation mutants ranging in length from 8 repeats to 26 repeats (Figure 1A). Spotting serial dilutions of actively growing yeast cultures then allowed us to score the growth of the truncated CTD mutants relative to wildtype controls.

**Figure 1.**
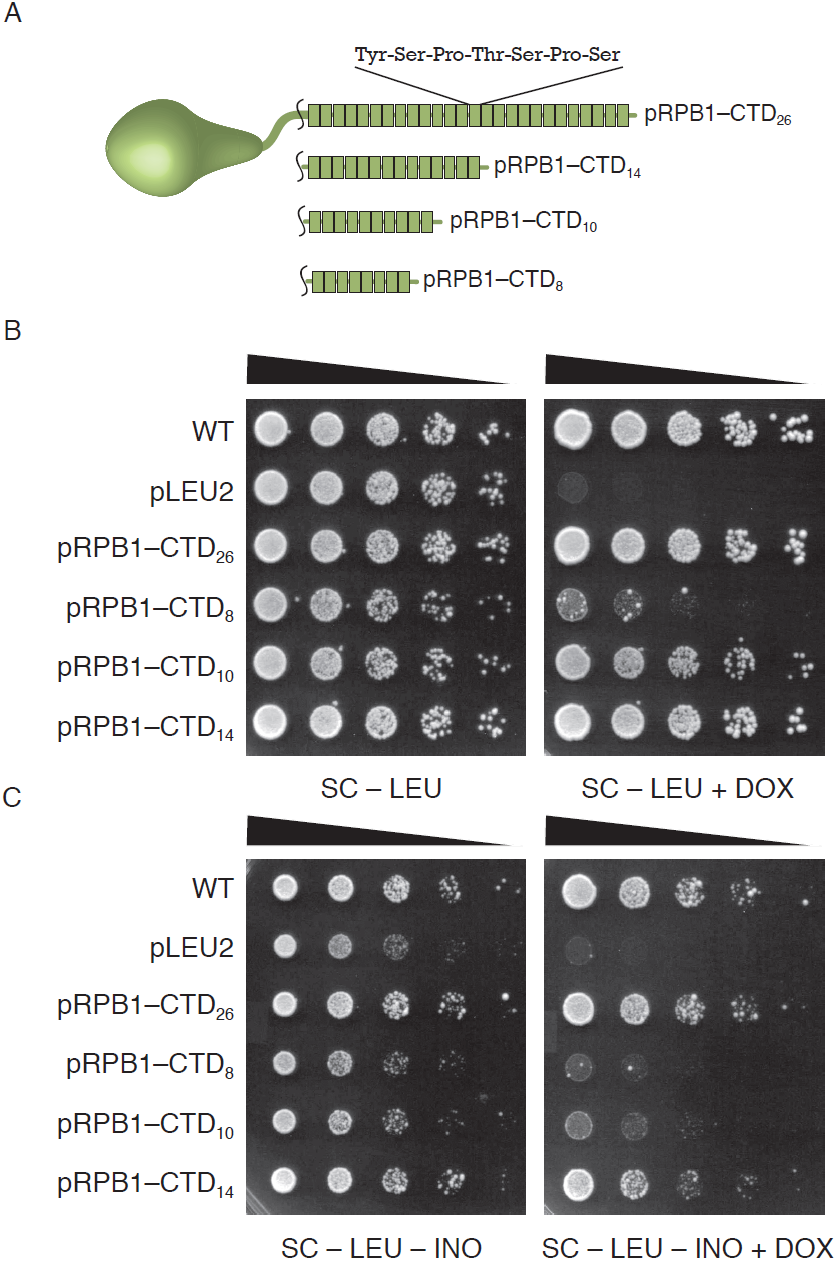
Length dependence of RNAPII CTD in TET-off system. (A) CTD truncation mutants tested in this study. Each block represents a single seven amino acid heptad repeat sequence. Constructs are labeled based on the number of total CTD repeats. (B) Spotting assay measuring the dependence of CTD length on yeast viability. In the absence of doxycycline (DOX) both the genomic copy of *RPB1* and the *LEU2* plasmid copy of *RPB1* harboring different length CTD regions are expressed. When DOX is present, only the plasmid copy is transcribed (MALAGON *et al.* 2006; MORRILL *et al.* 2016). (C) Spotting assay measuring the dependence of CTD length on yeast viability in media lacking inositol (INO).

In the absence of any stress, all cells grew equally well without DOX and this condition served as a loading control for our spotting assays. Strains labeled WT had the native budding yeast CTD sequence while the pRPB1-CTD_26_ construct had a synthetic full length CTD consisting of all perfect consensus repeats. Under all conditions tested the WT and CTD_26_ plasmids grew equally well and were considered equivalent. Addition of DOX led to a severe growth defect in the pRPB1–CTD_8_ construct, while the constructs with 10 and 14 repeats grew at wildtype levels (Figure 1B), in line with results from our previous work and the studies of other groups (NONET *et al.* 1987; MORRILL *et al.* 2016). We also assessed the ability of these truncation mutants to tolerate a number of nutrient and environmental stresses. pRPB1-CTD_10_ and pRPB1-CTD_14_ exhibited growth comparable to the full length CTD under all conditions except on media lacking inositol (Figure 1C and Figure S1). *INO1* is induced upon inositol starvation and both the CTD and its associated Mediator complex are required for this process (ARCHAMBAULT *et al.* 1996). The pRPB1–CTD_8_ mutant was inviable under the –INO+DOX condition while the pRPB1–CTD_10_ and pRPB1–CTD_14_ showed decreased growth relative to wildtype. In fact, even in the absence of DOX (when both the plasmid and genomic copies of RPB1 are expressed), the truncation plasmids have a slight effect on growth (compare Figure 1B left to Figure 1C left). Based on these observations we focused in on the inositol auxotrophy that results from mutating specific regions of the CTD.

### Positional requirements of CTD repeats in inositol auxotrophy

The graduated inositol auxotrophy of CTD mutants led to two hypotheses: growth on media lacking inositol requires either: 1) more than 8 CTD repeats, and approximately 14 repeats to achieve levels comparable to wild-type; or 2) CTD repeats in a particular linear position within the CTD sequence. Previous analysis of the CTD suggested that serine mutations in the consensus sequence had different effects if they were placed in proximal or distal repeats (WEST AND CORDEN 1995). We expanded on this work by creating a series of plasmids harboring serine to alanine (S>A) substitutions at different linear regions within the CTD (Figure 2A). The constructs each contained eight mutated repeats in a series of three windows while all maintaining 18 consensus repeats in various arrangements. The three block mutants were both expressed at similar levels and had comparable bulk Ser2 and Ser5 phosphorylation to the consensus plasmid pRPB1–CTD_26_ as measured by western blotting. Furthermore, overall Rpb1p levels in all constructs were reduced upon addition of DOX, reflecting the expected lack of expression from the DOX-regulated genomic copy of *RPB1* (Figure S2).

**Figure 2.**
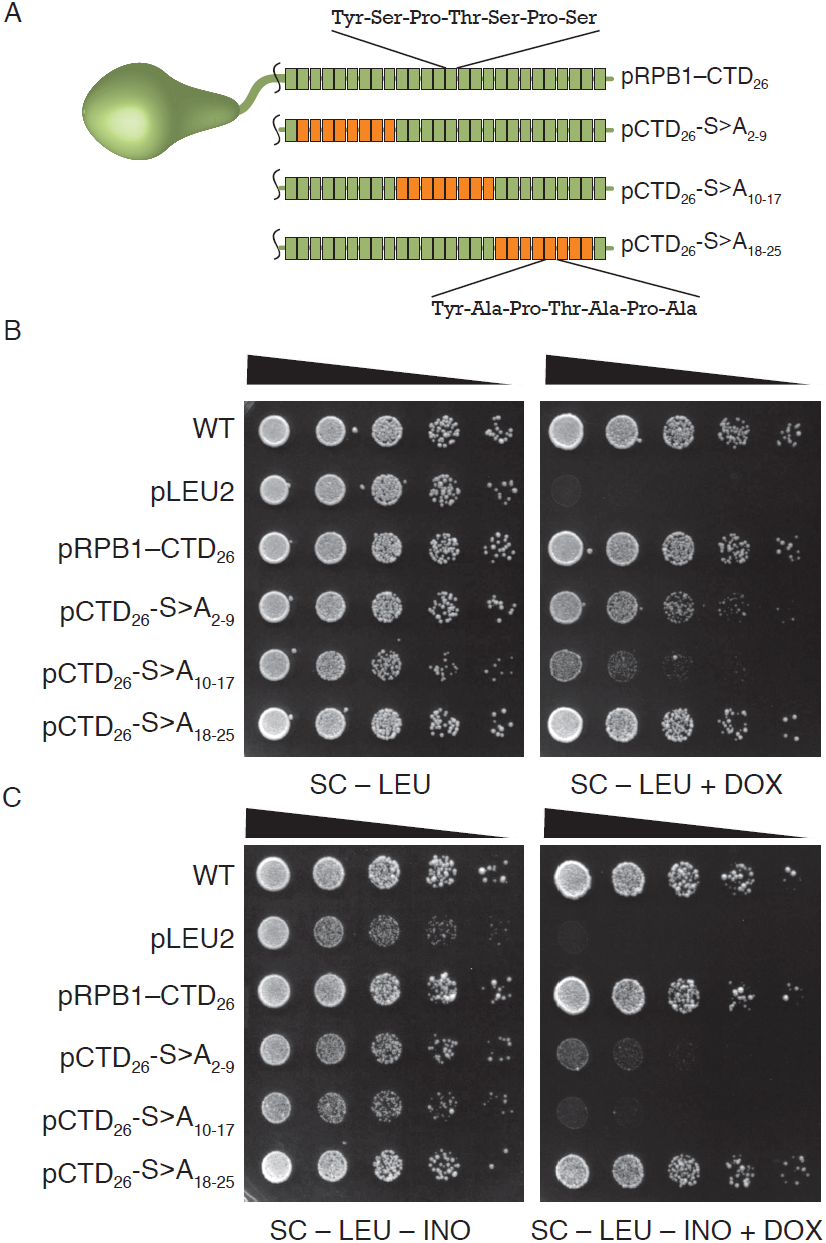
Position-specific phenotypes of CTD mutants. (A) CTD mutants were created that harbored Ser>Ala substitutions at precise positions within the CTD sequence as noted by the subscripts in the name. Repeats with wildtype sequence are colored in green with mutant repeats in orange. Spotting assay measuring the dependence of CTD position on yeast viability (B) and inositol auxotrophy (C).

In the absence of any stress, the pCTD_26_–S>A_18-25_ mutant behaved identically to the full-length consensus plasmid pRPB1–CTD_26_ (Figure 2B). Both pCTD_26_–S>A_2-9_ and pCTD_26_–S>A_10-17_ showed slower growth in the presence of DOX and in the absence of inositol (Figure 2B, 2C). In both the presence and absence of inositol, pCTD_26_–S>A_10-17_ was nearly inviable and we found this to be true for a number of additional phenotypic conditions as well. Specifically, the pCTD_26_–S>A_10-17_ mutant showed varying degrees of sensitivity to the drug 6-azauracil, galactose media and osmotic stress while the other two mutants were unaffected (Figure 3). However, for pCTD_26_–S>A_2-9_, while there was as slight growth defect on DOX, this mutant exhibited the same slow growth on media lacking inositol as seen for pRPB1–CTD_10_ and pRPB1–CTD_14_.

**Figure 3.**
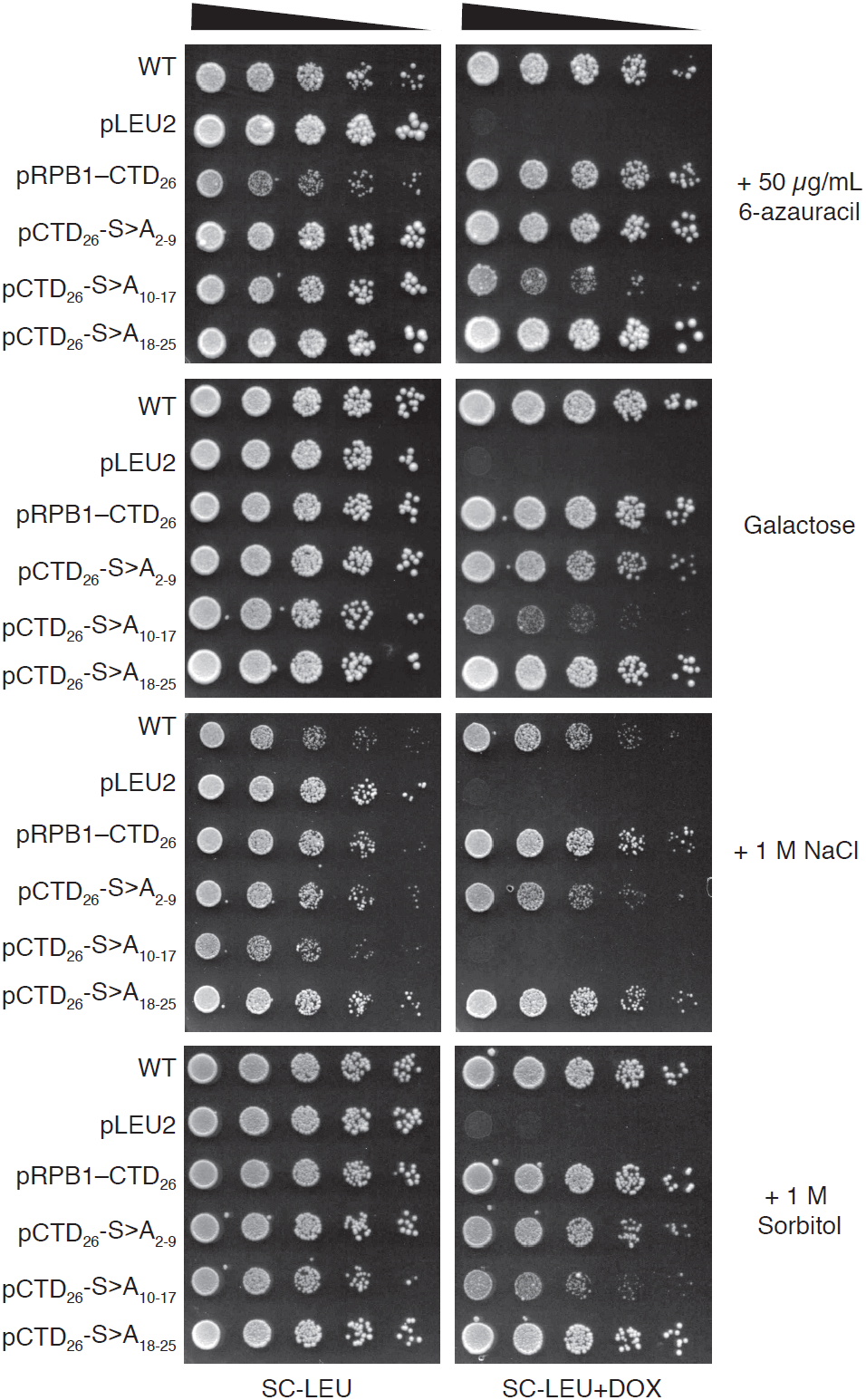
Additional phenotypes of position-specific CTD mutants. Preparation of the spotting assay and ordering of the mutants is the same as in Figure 2. CTD constructs were assayed on additional stresses including: 50 μg/mL of 6-Azauracil (6AU), plates with galactose as the only sugar (SC-GAL) and osmotic stress in the form of 1M NaCl and 1M sorbitol.

### Mutations in the CTD repeats 2–9 and 10–17 lead to impaired INO1 expression

The inositol auxotrophy phenotype has been extensively studied and mutations in the RNA polymerase II holoenzyme fail to induce expression at the *INO1* locus (ARCHAMBAULT *et al.* 1996). We suspected that poor growth of pCTD_26_–S>A_2-9_ and pCTD_26_–S>A_10-17_ on –INO+DOX was due to a similar lack of *INO1* induction. Therefore, we measured the induction of this gene in our positional mutants using endpoint RT-PCR with *ACT1* as a reference gene (Figure 4A). When strains were grown in the presence of inositol (+INO), *INO1* expression was completely suppressed while *ACT1* remained constant in both the –DOX and +DOX conditions. Under inducing conditions without DOX (–INO–DOX) all strains were able to induce *INO1*. Adding DOX to the inducing media (–INO+DOX) led to a sharp loss of *INO1* expression in both the pCTD_26_–S>A_2-9_ and the pCTD_26_–S>A_10-17_ mutants while *ACT1* expression remained constant. Representative gels are shown in Figure 4A while quantification of the RT-qPCR data for at least three independent cultures is shown in Figure 4B.

**Figure 4.**
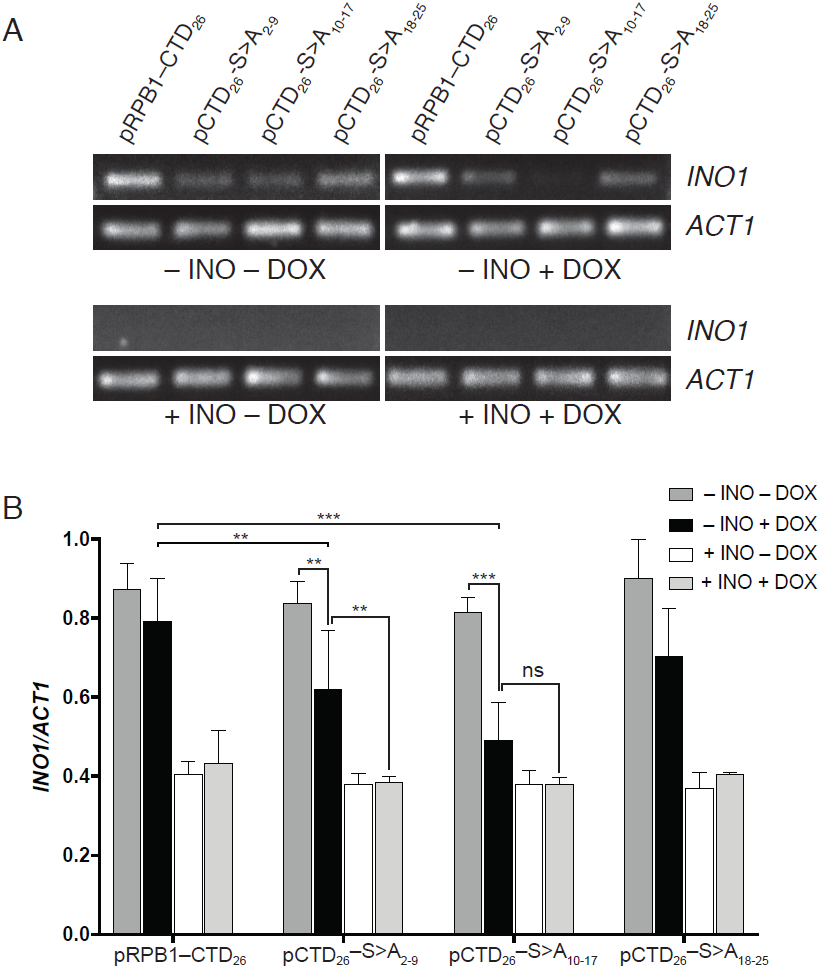
Effect of CTD position on *INO1* expression. (A) Representative agarose gels of RT-PCR reactions using primers specific for *INO1* and *ACT1* as a loading control. (B) Quantification of the effects of CTD mutation on *INO1* expression. Signal from agarose gels was quantified by densitometry using ImageJ and data are plotted as the ratio of the *INO1* band intensity to the *ACT1* band intensity. Two-way ANOVA was used to measure significance of interactions, and a subset of significant interactions are indicated as (**, adjusted P-value < 0.05; ***, adjusted P-value <0.01).

### Serine 5 is solely responsible for pCTD_26_–S>A_2-9_ inositol auxotrophy

The binding of protein factors to the CTD is determined, in part, by the modification state of defined residues in the heptad repeat (PHATNANI AND GREENLEAF 2006; WERNER-ALLEN *et al.* 2011). In order to determine which serine residue in the region-specific mutants contributed to the observed phenotypes we created a series of residue-specific mutants. These mutants have Ser2, Ser5 or Ser7 mutated to Ala within pCTD_26_–S>A_2-9_ (Figure 5A). Spotting the serine-specific mutants in the absence of stress led to wildtype levels of growth for all strains except pCTD_26_–S5A_2-9_ which showed the same slight growth defect as the original mutant (Figure 5B). Spotting the strains on –INO+DOX plates revealed that, indeed, pCTD_26_–S5A_2-9_ mirrored the slow growth phenotype of the pCTD_26_–S>A_2-9_ (Figure 5). Neither pCTD_26_–S2A_2-9_, nor pCTD_26_–S7A_2-9_, showed a growth defect without inositol despite having a similar number of serines mutated.Therefore, both the position of the serine with the heptad repeat, and its location within the linear CTD sequence is important for growth on media lacking inositol.

**Figure 5.**
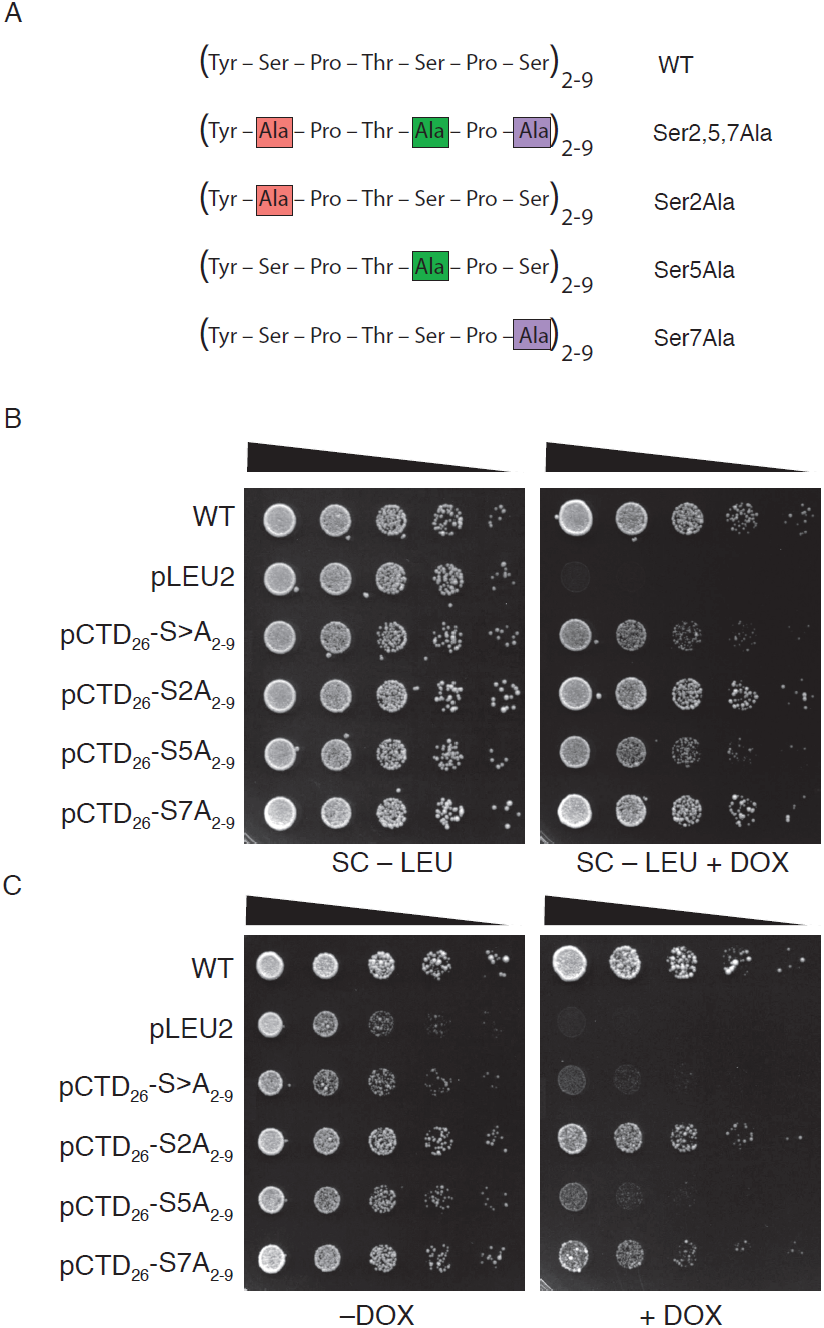
Influence of Ser>Ala substitutions on inositol auxotrophy in proximal CTD repeats. (A) CTD mutants expressing one or more Ser>Ala substitutions at discrete positions within repeats 2–9 of the RNAPII CTD. The position of the Ser>Ala substitution is marked in pink, noted in the name, and is carried by all 8 repeats within this region. Spotting assays measured the dependence of Ser position on yeast viability (B) and inositol auxotrophy (C).

### Suppressor mapping reveals two essential windows on the CTD

When plating pCTD_26_–S>A_2-9_ and pCTD_26_–S>A_10-17_ we observed that both yielded fast-growing suppressors when spotted on plates with DOX. Plasmid sequencing revealed two types of plasmid-based suppressors: homologous recombination with the genomic copy of *RPB1* or rearrangements of the repetitive CTD coding sequence itself to remove mutated repeats. We predicted that we could use the sequences of these suppressors to map important functional regions of the CTD. To screen for suppressors, 48 independent colonies of both the S>A_2-9_ and the S>A_10-17_ block mutants were grown overnight in a *rad52Δ* background, to bias towards rearrangements and away from homologous recombination with the genomic *RPB1* (MORRILL *et al.* 2016). Cells were spotted on +DOX and –INO+DOX plates and large, fast-growing colonies were isolated and analyzed by colony PCR and Sanger sequencing. Sequencing of over 30 independent contraction events revealed that the most common suppressors deleted either four or six mutant repeats or removed all variant repeats (resulting in truncations similar to Figure 1). Unexpectedly, we also recovered one suppressor of pCTD_26_–S>A_10-_ 17 which contained both a deletion and a duplication of variant repeats (Figure 6A and Figure S3).

**Figure 6.**
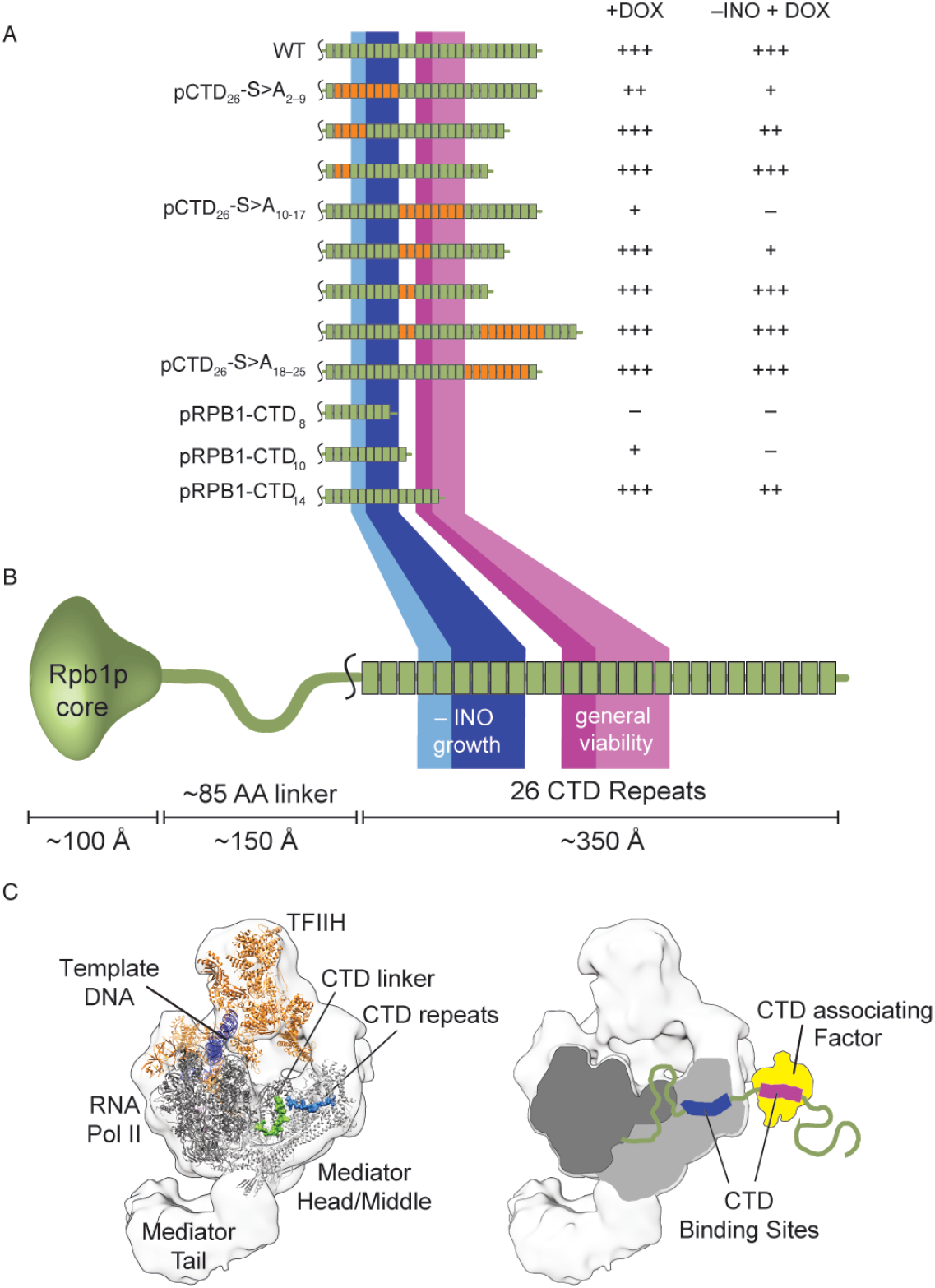
Mapping functional regions of the yeast CTD. (A) Summary of plasmid-based spontaneous suppressor mutants and their growth characteristics on general growth (+DOX) and inositol deficient (–INO+DOX) media. No growth is scored as (–) with poor, moderate, and unimpaired growth scored as +, ++, and +++, respectively. Representative spotting assays are available in Figure S4. (B) Based on the growth of different constructs in (A), Regions essential for general growth (purple), and important for growth in –INO media (blue) were mapped to the 26 repeats of the yeast CTD. Intensity of the color correlates with importance of a repeat for a particular phenotype. The scale bar represents approximate length of the CTD tail in a fully extended conformation, assuming a length of 2 Å/ amino acid for an extended peptide lacking secondary structure. (C) A model based on existing structures for the RNAPII in complex with Mediator proposing that Mediator interacts with repeats in the proximal region of the CTD, most likely repeats 5–9 (blue). This is spatially consistent with the binding of additional CTD associating factors (yellow) to the region defined as important for general viability (purple).

After confirming the identity of the suppressors, we isolated the plasmids and transformed them back into the original GRY3019 strain. These transformants were plated and their growth was scored on media containing DOX and/or inositol (Figure S3 and Figure 6A). Retransforming these plasmids confirmed that the improved growth is from a change in CTD sequence rather than another acquired mutation within the strain. Deleting four mutant repeats was sufficient to restore growth for both pCTD_26_–S>A_2-9_ and pCTD_26_–S>A_10-17_ when suppressors were spotted on +DOX plates. In contrast, deleting six mutant repeats from either block was necessary for a significant improvement in growth on –INO+DOX plates (Figure 6 and Figure S4). These repeat requirements enabled us to identify two repeat windows along the CTD that are necessary for viability and growth on media lacking inositol (Figure 6B).

### Structural modeling of CTD positional requirements

Our finding that the RNAPII CTD has two unique regions necessary for growth suggests that there exists at least two, non-overlapping binding sites for essential CTD-associating protein factors. Repeats 12-14 seem to be most important for growth and based on the strong phenotypes from mutations in this region it is difficult to predict which of the essential CTD-associated activities may be recruited to this site. However, both the importance of Ser5 and the inositol auxotrophy of pCTD_26_–S>A_2-9_ suggested to us that this region of the CTD may be important for recruitment of the Mediator complex. Deletion of some non-essential Mediator components leads to impaired growth on media lacking inositol (Figure S5). Thus to understand the geometric restraints of the CTD and gain insight into potential binding partners, we turned to recent cryo-electron microscopy (cryoEM) and x-ray crystal structures (ROBINSON *et al.* 2012; ROBINSON *et al.* 2016). Combining our suppressor data and current structural models, we built a to-scale representation of how the CTD tail might bind Mediator complex and a generic transcriptional regulator (Figure 6C). Based on the physical proximity between the core of Rpb1p and the Mediator-CTD binding site (only ∼50 Å), we propose that Mediator would most likely interact with the first CTD window, repeats 4-9.

## Discussion

The CTD of RNAPII is known to physically interact with a number of factors that are critical for cellular function including the capping enzyme complex, the Mediator complex, mRNA processing machinery, and termination factors such as Pcf11p (PHATNANI AND GREENLEAF 2006). It interacts with numerous additional non-essential factors such as the histone methyltransferase Set2p (KIZER *et al.* 2005). How protein factors organize on the RNAPII CTD during transcription has been of great interest for several decades. Early research dissected the importance of the heptad repeat sequence and the role of post-translational modifications toward the recruitment of factors (HSIN *et al.* 2011; SCHWER AND SHUMAN 2011). Mainly, the pattern of phosphorylation is known to change during different phases of transcription and this allows for the correct temporal recruitment of factors. Later mutational analysis revealed that the functional unit of the CTD consisted of two heptad repeats, with many protein factors binding the Ser5 region of the first repeat and the Ser2 region of the next (STILLER AND COOK 2004; LIU *et al.* 2008). Due to the repetitive nature of the CTD, it has been difficult to determine whether different repeats within the linear sequence had specific functions. Early synthetic mutants showed variable phenotypes depending on the location of the mutation but these mutants were neither systematic nor uniform in length (WEST AND CORDEN 1995). In mammalian cells, the binding of certain factors has been reported to be specific to either the N or C-terminal halves of the repeats (FONG AND BENTLEY 2001; YOH *et al.* 2008). However, the seventh residue in the C-terminal repeats is frequently degenerate, therefore both sequence and spatial positioning may determine function. Currently, we collectively know a lot about how different factors can recognize a region of the CTD but there is still little known about how they might simultaneously interact with the full-length CTD. For example, are all repeats functionally equivalent, or is there undescribed specificity built into the linear CTD sequence? Additionally, is factor binding controlled by both phosphorylation state and competition with other factors? Or are there higher-order structural interactions that provide targeting of particular factors to specific regions of the CTD.

We recently reported on an improved TET-off system to investigate CTD mutants (MORRILL *et al.* 2016). With this system, it became possible to make precise mutations to different regions of the CTD. Here, we utilized our TET-off system to explore whether specific repeats had essential CTD functions. We constructed three different block mutants containing Ser to Ala mutations in the heptad repeats and uncovered different effects for all three regions. If all the CTD repeats had identical function as implied by their primary amino acid sequence, then we would expect all three of our block mutants to behave equally. Indeed, our block mutants had similar expression levels and serine phosphorylation profiles to each other and to the pRPB1–CTD_26_ control (Figure S2). However, we found that all three blocks still behaved differently. Critically, while all three mutants contained 18 wildtype repeats in various arrangements (Figure 2A) the first two blocks yielded different phenotypes. Given that 14 wildtype repeats permit growth under all tested conditions (Figure 1B, C), we can rule out bulk repeat number as the only determinant for CTD function. Instead we propose that our mutations are interrupting positional cues within the CTD sequence that coordinate the binding of protein factors. We examined a number of different phenotypes using our block mutants and found that mutating repeats 10–17 caused a general sensitivity to stress conditions. The sensitivity was comparable to that of the pRPB1–CTD_8_ mutant and even led to complete inviability in the presence of 1M NaCl (Figure 3). One result from our screen that stood out in particular was the unique sensitivity of the pCTD_26_–S>A_2-9_ mutant to the inositol auxotrophy phenotype (Figure 2C). Inositol auxotrophy has been a commonly observed phenotype in transcription mutants (VILLA-GARCIA *et al.* 2011). In particular, mutations in both the CTD and the Mediator complex have been shown to cause inositol auxotrophy (ARCHAMBAULT *et al.* 1996; SINGH *et al.* 2006). Although disruptions in other physiological pathways can lead to inositol auxotrophy (YOUNG *et al.* 2010), our pCTD_26_–S>A_2-9_ and pCTD_26_–S>A_10-17_ mutants fail to express *INO1*, demonstrating that these regions are required for inducible transcription (Figure 4). Based on the specific inositol auxotrophy when repeats 2–9 were mutated, we further probed this region using a set of residue-specific mutants. Mutating Ser5 within repeats 2–9 resulted in the same inositol auxotrophy observed when all three serine residues within these repeats were replaced with alanine (Figure 5B). This requirement for Ser5 is consistent with a number of known CTD binding proteins including Kin28 and the Mediator complex (ROBINSON *et al.* 2012; WONG *et al.* 2014) as well as the 5’ capping enzyme (FABREGA *et al.* 2003). An analysis of the spontaneous suppressors we found allowed us to further define the regions of the CTD important for function. Mapping these suppressors identified two regions, repeats 6–9 and 14–17, that were required for growth on +DOX plates. Most CTD-binding factors use two repeats to bind while the largest known interaction is with three repeats and the Mediator complex (KUBICEK *et al.* 2012; ROBINSON *et al.* 2012). Therefore, it is highly unlikely that both regions of four repeats each are bound by a single protein factor. Additionally, the different phenotypes observed when repeats 2–9 or 10–17 are mutated (Figure 2B, C) support at least two independent binding events that are required at repeats 6–9 and 14–17. These two regions expand to require repeats 4–9 and 12–17 for growth on media lacking inositol.

Intriguingly, repeats 10 and 11, which reside between the two CTD regions defined in this work, are dispensable for growth under all tested conditions. Previous mutational analysis of the CTD demonstrated that spacers of two or five Ala residues could still maintain viability provided they were inserted between every functional diheptad (STILLER AND COOK 2004). Consequently, non-functional sequences are tolerated provided the spacing of essential repeats is not disrupted. Thus, repeats 10 and 11 may be acting as natural structural spacers that help align the essential regions in repeats 4– 9 and 12–17 for separate binding events.

To address the possibilities raised by our genetic data we attempted to use existing RNAPII and Mediator structural data to model how Mediator may interact with specific CTD repeats. The best structural evidence available suggests that Rpb1p and the known CTD-Mediator binding site are only 50 Å apart. While the length of the linker would allow Mediator to bind any CTD repeat, we propose that CTD repeats 4–9 are the primary site of Mediator association. This arrangement is consistent with the requirement of more than eight CTD repeats for viability (NONET *et al.* 1987; WEST AND CORDEN 1995; MORRILL *et al.* 2016) and more than 12 repeats for normal growth. If the Mediator binding site is confined to the 4–9 window, this would allow a second binding event within the 13-CTD tail, and could explain why further truncations are inviable. Mediator interactions require an unphosphorylated Ser5 within the CTD for binding (JERONIMO AND ROBERT 2014), consistent with our pCTD_26_–S5A_2-9_ mutant under inositol auxotrophy (Figure 5C). Mediator is required for growth without inositol and a number of nonessential Mediator subunits demonstrate inositol auxotrophy when deleted (SINGH *et al.* 2006; YOUNG *et al.* 2010) and (Figure S5). Although we found that both the 4–9 and 12–17 windows are required to survive without inositol (Figure 6A), the second window is also required for viability under a wide range of stresses (Figure 2C, Figure 3). Thus, we reason that the 4–9 window harbors an exclusive Mediator binding site while the 12– 17 window is used for other essential CTD-related pathways.

The model we present shows one conformation of Mediator and other factors bound to the two essential regions we identified in our genetic screen. However, the dynamic nature of both the CTD (ZHANG *et al.* 2010) and the Mediator complex (SENNETT AND TAATJES 2014; WANG *et al.* 2014) means that alternative binding conformations are also possible. Residues as far away as the very tip of the CTD are able to make contacts with Mediator subunits (NOZAWA *et al.* 2017), raising the prospect that either of our two windows might bind to Mediator. Mapping suppressors revealed that repeats 12–17 are also required for viability under inositol auxotrophy (Figure 6A) and may represent a possible binding site for Mediator. The Mediator complex, or some other factor required for *INO1* induction, may either bind this 12–17 window or possibly sample both the 4–9 and 12–17 windows to properly coordinate transcription. Interestingly, while many deletion mutants of non-essential Mediator subunits demonstrate inositol auxotrophy (SINGH *et al.* 2006; YOUNG *et al.* 2010), the severity of the defect varies based on the subunit deleted. We found that while some mutants (e.g. *srb5Δ, rox3Δ*) were inviable when grown without inositol, other mutants (e.g. *srb2Δ, soh1Δ*) showed only a reduced growth rate (Figure S5). These growth differences recall the different phenotypes of our pCTD_26_–S>A_2-9_ and pCTD_26_–S>A_10-17_ mutants and raise the possibility that Mediator complexes or subcomplexes of differing subunit composition may selectively bind either of the two essential CTD windows.

In addition to the Mediator complex, there are a number of other possible CTD-binding proteins that may specifically occupy the CTD windows at repeats 4–9 and 12–17. Similarly to Mediator subunits, CTD kinases Ssn3p and Ctk1p have been identified in screens for inositol auxotrophy (YOUNG *et al.* 2010). Our Ser to Ala substitution mutants in the 4–9 window could be preventing proper phosphorylation at these repeats even if Mediator is productively bound at the 12–17 window. While we did not detect any differences in Ser2 and Ser5 phosphorylation in our CTD mutants (Figure S2), a region-specific defect in phosphorylation may be too subtle to be detected or restricted to a certain stress condition. Alternatively, other essential co-transcriptional processes such as mRNA 5’ capping have also been shown to require properly modified Ser5 (FABREGA *et al.* 2003) and capping enzyme may bind to repeats 4–9 while mediator occupies the 12–17 site. The elongation factor Spt4p has also been associated with the inositol auxotrophy phenotype (YOUNG *et al.* 2010) and could potentially bind to one of the two CTD windows in a sequential manner following Mediator or a CTD kinase.

Discriminating between these multiple possible binding configurations will require biochemical characterization of the RNA polymerase II complexes across our various region-specific CTD constructs.

Most fundamentally, the analysis here demonstrates conclusively that, although they have the same amino acid sequence, different heptads of the CTD have specific cellular functions. Repeats 12–17 are important for growth on a range of phenotypes whereas repeats 4–9 are required specifically for growth in the absence of inositol which is consistent with these repeats being important for Mediator binding. This solidifies CTD repeat location, in addition to CTD phosphorylation, as an important factor in determining how CTD-associating proteins interact with the CTD during transcription. Using our approach and growing panel of site-specific mutants, it should be possible to probe even more specific CTD interactions with factors ranging from the RNA capping and processing machinery to chromatin modifying enzymes.

---

Figure S1. Additional phenotypes of CTD truncation mutants. Preparation of the spotting assay and ordering of the mutants is the same as in Figure 1. Conditions tested are: 37° C, 50 μg/mL of 6-Azauracil (6AU), plates with galactose as the only sugar (SC-GAL) and 1M urea. Pictures were taken starting at two days and a representative image of three independent experiments is presented.

Figure S2. Expression and phosphorylation levels of position-specific mutants. Western blot of CTD constructs grown in the presence and absence of DOX. Total Rpb1p levels as well as Ser2 and Ser5 phosphorylation were assayed and compared to a G6PDH housekeeping gene loading control. A long exposure panel is provided for the Rpb1p and Ser2phos blots due to the faint signal observed for the +DOX samples.

Figure S3. Sequence alignments of CTD coding region from pCTD_26_–S>A_2-9_ and pCTD_26_–S>A_10-17_ and corresponding suppressor mutants. Changes in CTD length that were observed by colony PCR were confirmed by Sanger sequencing. The position of mutated repeats is highlighted in pink. (A) Alignment of suppressors with either four (Δ4) or six (Δ6) mutant repeats deleted from the pCTD_26_–S>A_2-9_ region-specific mutant. (B) Alignment of suppressors with either four (Δ4) or six (Δ6) mutant repeats deleted or a more complex rearrangement (Δ12-21^(2-17)) from the pCTD_26_–S>A_10-17_ mutant.

Figure S4. Improved growth of region-specific suppressors. Suppressor plasmids were retransformed into the tet-off strain GRY3019 and scored for growth on standard (+DOX) and inositol deficient (–INO+DOX) media. (A) Spotting assay for suppressor mutants of the pCTD_26_–S>A_2-9_ region-specific mutant. (B) Spotting assay for suppressor mutants of the pCTD_26_–S>A_10-17_ region-specific mutant.

Figure S5. Phenotypes of Mediator subunit deletion strains. Yeast strains harboring deletions of the listed Mediator subunits were grown up and spotted on the indicated plate types. Strains were grouped based upon their predicted localization into the Cdk8, head, middle or tail subcomplexes (MALIK AND ROEDER 2010)

## References

Adams, A., D. E. Gottschling, C. A. Kaiser and T. Stearns, 1997 Methods in Yeast Genetics: A Cold Spring Harbor Laboratory Course Manual. Cold Spring Harbor Press, Plainview, NY.

Archambault, J., D. B. Jansma and J. D. Friesen, 1996 Underproduction of the largest subunit of RNA polymerase II causes temperature sensitivity, slow growth, and inositol auxotrophy in Saccharomyces cerevisiae. Genetics 142: 737–747.

Bartolomei, M. S., N. F. Halden, C. R. Cullen and J. L. Corden, 1988 Genetic analysis of the repetitive carboxyl-terminal domain of the largest subunit of mouse RNA polymerase II. Mol Cell Biol 8: 330–339.

Corden, J. L., 2013 RNA polymerase II C-terminal domain: Tethering transcription to transcript and template. Chem Rev 113: 8423–8455.

Eick, D., and M. Geyer, 2013 The RNA polymerase II carboxy-terminal domain (CTD) code. Chem Rev 113: 8456–8490.

Feaver, W. J., O. Gileadi, Y. Li and R. D. Kornberg, 1991 CTD kinase associated with yeast RNA polymerase II initiation factor b. Cell 67: 1223–1230.

Fong, N., and D. L. Bentley, 2001 Capping, splicing, and 3’ processing are independently stimulated by RNA polymerase II: different functions for different segments of the CTD. Genes Dev 15: 1783–1795.

Fuchs, S. M., R. N. Laribee and B. D. Strahl, 2009 Protein modifications in transcription elongation. Biochim Biophys Acta 1789: 26–36.

Greber, B. J., T. H. D. Nguyen, J. Fang, P. V. Afonine, P. D. Adams et al., 2017 The cryo-electron microscopy structure of human transcription factor IIH. Nature 549: 414–417.

Hsin, J. P., A. Sheth and J. L. Manley, 2011 RNAP II CTD phosphorylated on threonine-4 is required for histone mRNA 3’ end processing. Science 334: 683–686.

Janke, C., M. M. Magiera, N. Rathfelder, C. Taxis, S. Reber et al., 2004 A versatile toolbox for PCR-based tagging of yeast genes: new fluorescent proteins, more markers and promoter substitution cassettes. Yeast 21: 947–962.

Kubicek, K., H. Cerna, P. Holub, J. Pasulka, D. Hrossova et al., 2012 Serine phosphorylation and proline isomerization in RNAP II CTD control recruitment of Nrd1. Genes Dev 26: 1891–1896.

Lee, J. M., and A. L. Greenleaf, 1989 A protein kinase that phosphorylates the C-terminal repeat domain of the largest subunit of RNA polymerase II. Proc Natl Acad Sci U S A 86: 3624–3628.

Liu, P., A. L. Greenleaf and J. W. Stiller, 2008 The essential sequence elements required for RNAP II carboxyl-terminal domain function in yeast and their evolutionary conservation. Mol Biol Evol 25: 719–727.

Malagon, F., M. L. Kireeva, B. K. Shafer, L. Lubkowska, M. Kashlev et al., 2006 Mutations in the Saccharomyces cerevisiae RPB1 gene conferring hypersensitivity to 6-azauracil. Genetics 172: 2201–2209.

McDaniel, J. R., J. A. Mackay, F. G. Quiroz and A. Chilkoti, 2010 Recursive directional ligation by plasmid reconstruction allows rapid and seamless cloning of oligomeric genes. Biomacromolecules 11: 944–952.

Morrill, S. A., A. E. Exner, M. Babokhov, B. I. Reinfeld and S. M. Fuchs, 2016 DNA Instability Maintains the Repeat Length of the Yeast RNA Polymerase II C-terminal Domain. J Biol Chem 291: 11540–11550.

Nonet, M., D. Sweetser and R. A. Young, 1987 Functional redundancy and structural polymorphism in the large subunit of RNA polymerase II. Cell 50: 909–915.

Nonet, M. L., and R. A. Young, 1989 Intragenic and extragenic suppressors of mutations in the heptapeptide repeat domain of Saccharomyces cerevisiae RNA polymerase II. Genetics 123: 715–724.

Pettersen, E. F., T. D. Goddard, C. C. Huang, G. S. Couch, D. M. Greenblatt et al., 2004 UCSF Chimera--a visualization system for exploratory research and analysis. J Comput Chem 25: 1605–1612.

Phatnani, H. P., and A. L. Greenleaf, 2006 Phosphorylation and functions of the RNA polymerase II CTD. Genes Dev 20: 2922–2936.

Powell, W., and D. Reines, 1996 Mutations in the second largest subunit of RNA polymerase II cause 6-azauracil sensitivity in yeast and increased transcriptional arrest in vitro. J Biol Chem 271: 6866–6873.

Robinson, P. J., D. A. Bushnell, M. J. Trnka, A. L. Burlingame and R. D. Kornberg, 2012 Structure of the mediator head module bound to the carboxy-terminal domain of RNA polymerase II. Proc Natl Acad Sci U S A 109: 17931–17935.

Robinson, P. J., M. J. Trnka, D. A. Bushnell, R. E. Davis, P. J. Mattei et al., 2016 Structure of a Complete Mediator-RNA Polymerase II Pre-Initiation Complex. Cell 166: 1411–1422 e1416.

Schneider, S., Y. Pei, S. Shuman and B. Schwer, 2010 Separable functions of the fission yeast Spt5 carboxyl-terminal domain (CTD) in capping enzyme binding and transcription elongation overlap with those of the RNA polymerase II CTD. Mol Cell Biol 30: 2353–2364.

Schwer, B., and S. Shuman, 2011 Deciphering the RNA polymerase II CTD code in fission yeast. Mol Cell 43: 311–318.

Spahr, H., G. Calero, D. A. Bushnell and R. D. Kornberg, 2009 Schizosacharomyces pombe RNA polymerase II at 3.6-A resolution. Proc Natl Acad Sci U S A 106: 9185–9190.

Stiller, J. W., and M. S. Cook, 2004 Functional unit of the RNA polymerase II C-terminal domain lies within heptapeptide pairs. Eukaryot Cell 3: 735–740.

Suh, H., S. B. Ficarro, U. B. Kang, Y. Chun, J. A. Marto et al., 2016 Direct Analysis of Phosphorylation Sites on the Rpb1 C-Terminal Domain of RNA Polymerase II. Mol Cell 61: 297–304.

Valay, J. G., M. Simon, M. F. Dubois, O. Bensaude, C. Facca et al., 1995 The KIN28 gene is required both for RNA polymerase II mediated transcription and phosphorylation of the Rpb1p CTD. J Mol Biol 249: 535–544.

Villa-Garcia, M. J., M. S. Choi, F. I. Hinz, M. L. Gaspar, S. A. Jesch et al., 2011 Genome-wide screen for inositol auxotrophy in Saccharomyces cerevisiae implicates lipid metabolism in stress response signaling. Mol Genet Genomics 285: 125–149.

Werner-Allen, J. W., C. J. Lee, P. Liu, N. I. Nicely, S. Wang et al., 2011 cis-Proline-mediated Ser(P)5 dephosphorylation by the RNA polymerase II C-terminal domain phosphatase Ssu72. J Biol Chem 286: 5717–5726.

West, M. L., and J. L. Corden, 1995 Construction and analysis of yeast RNA polymerase II CTD deletion and substitution mutations. Genetics 140: 1223–1233.

Wilcox, C. B., A. Rossettini and S. D. Hanes, 2004 Genetic interactions with C-terminal domain (CTD) kinases and the CTD of RNA Pol II suggest a role for ESS1 in transcription initiation and elongation in Saccharomyces cerevisiae. Genetics 167: 93–105.

Wong, K. H., Y. Jin and K. Struhl, 2014 TFIIH phosphorylation of the Pol II CTD stimulates mediator dissociation from the preinitiation complex and promoter escape. Mol Cell 54: 601–612.

Yoh, S. M., J. S. Lucas and K. A. Jones, 2008 The Iws1:Spt6:CTD complex controls cotranscriptional mRNA biosynthesis and HYPB/Setd2-mediated histone H3K36 methylation. Genes Dev 22: 3422–3434.

Zhang, J., and J. L. Corden, 1991 Phosphorylation causes a conformational change in the carboxyl-terminal domain of the mouse RNA polymerase II largest subunit. J Biol Chem 266: 2297–2302.

